# A systematic evaluation of interneuron morphology representations for cell type discrimination

**DOI:** 10.1101/591370

**Authors:** Sophie Laturnus, Dmitry Kobak, Philipp Berens

## Abstract

Quantitative analysis of neuronal morphologies usually begins with choosing a particular feature representation in order to make individual morphologies amenable to standard statistics tools and machine learning algorithms. Many different feature representations have been suggested in the literature, ranging from density maps to intersection profiles, but they have never been compared side by side. Here we performed a systematic comparison of various representations, measuring how well they were able to capture the difference between known morphological cell types. For our benchmarking effort, we used several curated data sets consisting of mouse retinal bipolar cells and cortical inhibitory neurons. We found that the best performing feature representations were two-dimensional density maps closely followed by morphometric statistics, which both continued to perform well even when neurons were only partially traced. The same representations performed well in an unsupervised setting, implying that they can be suitable for dimensionality reduction or clustering.

## 1 Introduction

The development of experimental methods for high-throughput single cell RNA sequencing (Zeisel et al., 2018; Saunders et al., 2018; Tasic et al., 2018; Cao et al., 2019) and large-scale functional imaging (Baden et al., 2016; Pachitariu et al., 2017; Schultz et al., 2017) has led to a surge of interest in identifying the building blocks of the brain – the neural cell types (Zeng and Sanes, 2017; Chen et al., 2018). Both data modalities are analyzed with specialized quantitative tools (Stegle et al., 2015; Stringer and Pachitariu, 2019) and produce data sets amenable to statistical analysis such as cell type identification by clustering.

At the same time, ever since the work of Santiago Ramón y Cajal (Ramón y Cajal, 1899), it was the anatomy of a neuron that has been considered the defining feature of a neural cell type. Like in genetics and physiology, recent years have seen a tremendous increase in the availability of anatomical data sets, due to advances in light and electron microscopy (Briggman et al., 2011; Helmstaedter et al., 2013; Economo et al., 2016) and associated tools for increasingly automated reconstruction (Peng et al., 2010, 2014; Bria et al., 2016). As a consequence, more and more full reconstructions of neurons are becoming available in public databases, such as the Allen cell type atlas (http://celltypes.brain-map.org) or the NeuroMorpho database (http://neuromorpho.org).

Anatomical analysis of neural cell types based on these reconstructions, however, requires accurate quantitative representations of the neuron morphologies. While many different representations have been developed in the literature, they have rarely been systematically compared with regard to their ability to discriminate different cell types. Two prominent examples of such representations are density maps (Jefferis et al., 2007) and morphometric statistics (Uylings and van Pelt, 2002; Scorcioni et al., 2008; Polavaram et al., 2014; Lu et al., 2013, 2015), representing two ends of the spectrum: density maps ignore all fine details of a morphology, simply measuring the density of neurites; morphometric statistics, in turn, quantify the complex branching of axons and dendrites in a set of single-valued summary statistics. Other spatial analyses such as Sholl intersection profiles (Sholl, 1953) can be seen as occupying an intermediate position on this spectrum. In addition, several novel feature representations based on graph theory and topology have been suggested in recent years (Heumann and Wittum, 2009; Gillette and Grefenstette, 2009; Gillette et al., 2015; Li et al., 2017; Kanari et al., 2018).

Here we benchmarked different representations of neural morphologies as to how well they were able to capture the difference between known morphological types of interneurons. We used carefully curated anatomical data from three studies, encompassing over 500 retinal and cortical interneurons with complete axonal and dendritic reconstructions and expert annotated cell type labels (Helmstaedter et al., 2013; Jiang et al., 2015; Scala et al., 2019). In order to have a well-defined performance measure, we used a supervised learning framework: given the expert labels, we asked which morphological representations were most suitable for cell type discrimination. By combining different representations together, we also studied to what extent they captured complementary information about cell morphologies. In addition, we investigated how robust these representations are if only parts of a neuron are reconstructed and how useful they remain in an unsupervised setting.

## 2 Results

### 2.1 Morphological feature representations

We analyzed the discriminability between different morphological cell types in adult mouse retina and adult mouse cortex (Figure 1). The retinal data set consisted of *n* = 221 retinal bipolar cells semi-automatically reconstructed from electron microscopy scans and sorted into 13 distinct cell types (Helmstaedter et al., 2013; Behrens et al., 2016) (Figure 1A). In this study we only used the 11 cell types that included more than 5 neurons (remaining sample size *n* = 212). The cortical data consisted of inhibitory interneurons from primary visual cortex manually reconstructed based on biocytin stainings (Jiang et al., 2015; Scala et al., 2019). We analyzed the neurons separated by layer (V1 L2/3: *n* = 108 neurons in 7 classes, Figure 1B; V1 L4: *n* = 92 neurons in 7 classes, Figure 1C; V1 L5: *n* = 93 neurons in 6 classes, Figure 1D). All four data sets comprised accurate and complete morphological reconstructions of dendrites and axons and included cell types that are morphologically close enough to pose a challenge for classification (see Discussion).

**Figure 1:**
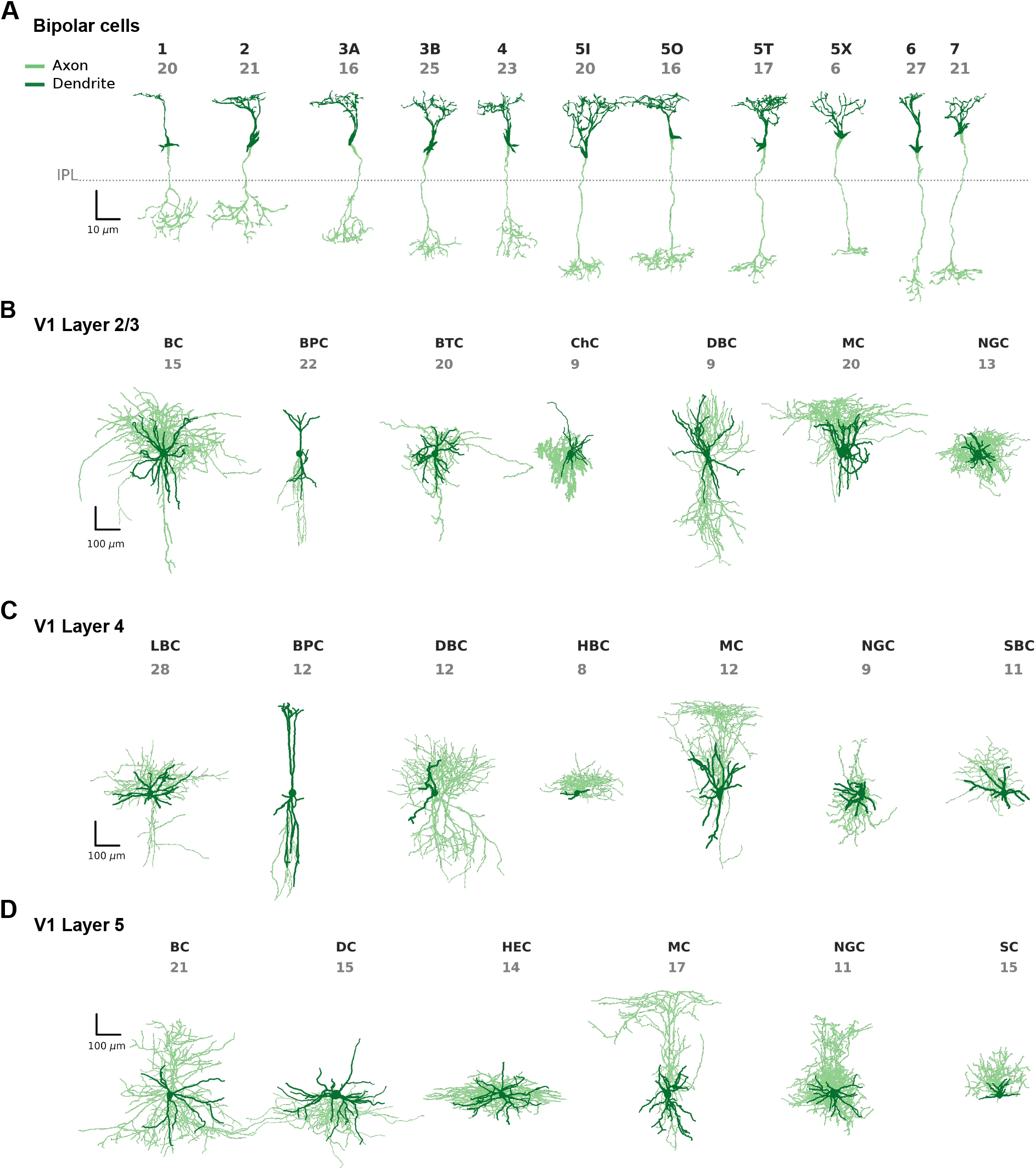
Exemplary cells of each cell type for all five data sets. Axons are shown in light green, dendrites in dark green. **(A)**Mouse retinal bipolar cells (cone-connecting) from Helmstaedter et al. (2013). The dashed line shows the onset of the inner plexiform layer (IPL). The cell types used for analysis are types 1, 2, 3A, 3B, 4, 5I, 5O, 5T, X, 6, and 7. Cell types 8 and 9 were excluded from further analysis due to insufficient sample sizes. **(B)**Layer 2/3 inhibitory interneurons in primary visual cortex of adult mice (Jiang et al., 2015). BC: basket cells, BPC: bipolar cells, BTC: bitufted cells, ChC: chandelier cells, DBC: double bouquet cells, MC: Martinotti cells, NGC: neurogliaform cells. **(C)**Layer 4 inhibitory interneurons in primary visual cortex of adult mice (Scala et al., 2019). LBC: large basket cells, BPC: bipolar cells, DBC: double bouquet cells, HBC: horizontal basket cells, MC: Martinotti cells, NGC: neurogliaform cells, SBC: small basket cells. **(D)**Layer 5 inhibitory interneurons in primary visual cortex of adult mice (Jiang et al., 2015). BC: basket cells, DC: deep-projecting cells, HEC: horizontally elongated cells, MC: Martinotti cells, NGC: neurogliaform cells, SC: shrub cells.

We investigated 62 feature representations that we grouped into four different categories: density maps, morphometric statistics, morphometric distributions, and persistence images (Figure 2). Each feature representation was computed using only axons, only dendrites, and using the full neuron (i.e. axons and dendrites together).

**Figure 2:**
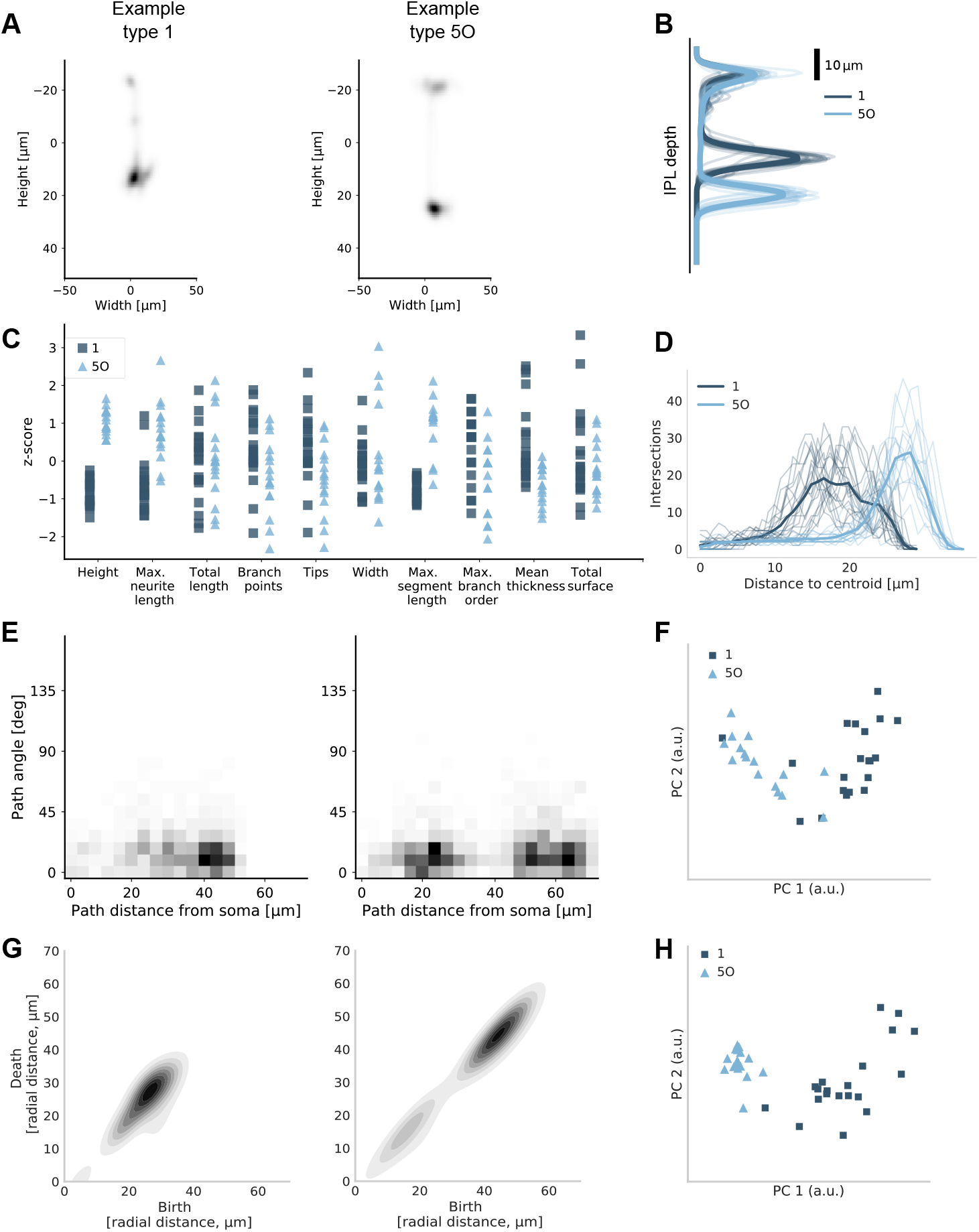
Selected feature representations for retinal bipolar cells of type 1 and type 5O. **(A)** Smoothed density map of XZ projection for two exemplary cells. **(B)** Smoothed density map of Z projection for all cells of these two types. The cells of type 5O stratify deeper in the inner plexiform layer (IPL) than cells of type 1. Bold lines show class means. **(C)** A selection of ten single-valued summary statistics that were included in the *morphometric statistics* vector. **(D)** Sholl intersection profile of the YZ projection for all cells of these two types. Bold lines show class means. **(E)** Two-dimensional distribution of path angles and path distances to the soma across all nodes for the same two exemplary cells shown in (A). **(F)** The first and the second principal components (PCs) of path-angle/path-distance histograms for all cells of these two types. **(G)** Two-dimensional persistence images for the same two exemplary cells shown in (A) and (E). **(H)** The first and the second PCs of 2D persistence images for all cells of these two types.

Density maps are one- or two-dimensional projections of the neural morphology. We used projections onto the *x*, *y*, and *z* axes as well as onto the *xy*, *xz*, and *yz* planes. Figure 2A shows the XZ density maps for two exemplary bipolar cells, one of type 1 and one of type 5O. Figure 2B shows Z density maps of all cells of these two types. This particular pair of cell types can be easily discriminated based on the Z projection alone.

We used 24 single-valued summary statistics of each neuron, such as width, height, total neurite length, number of tips, number of branch points, etc., many of which were different for the bipolar types 1 and 5O (Figure 2C). We also considered a feature representation that joins all of them into a 24-dimensional morphometric statistics vector.

We used 23 morphometric distributions of which 17 were one-dimensional and six were two-dimensional. As an example, the Sholl intersection profile (Sholl, 1953) describes the number of intersection of a 2D projection with concentric circles of different radius, and is very different for bipolar types 1 and 5O. (Figure 2D). An example of a two-dimensional distribution is the distribution of path angle (turning angle) vs. path distance (distance to soma along the neurite path) across all nodes in the traced morphology (Figure 2E). After binning, this becomes a 400-dimensional feature vector; Figure 2F shows two principal components (PCs) across all bipolar cells of type 1 and 5O, indicating that PC1 discriminates the types very well.

Finally, we used persistence images, a recently introduced quantification of neural morphology based on topological ideas (Li et al., 2017; Kanari et al., 2018, 2019). We used four different distance functions (also called filter functions) to construct one- and two-dimensional persistence images, resulting in eight different persistence representations. The same two bipolar cell types can be well discriminated based on PC1 of the two-dimensional radial-distance-based persistence images (Figure 2G–H).

See Methods for a complete list and detailed definitions of the investigated feature representations.

### 2.2 Predictive performance of feature representations

For each feature representation and for each pair of morphological types in a given data set, we built a binary classifier and assessed its performance using cross-validation. As a classifier, we used logistic regression regularized with elastic net penalty and PCA pre-processing. Nested cross-validation was used to tune the regularization strength and obtain an unbiased estimate of the performance (see Methods and Figure 3).

**Figure 3:**
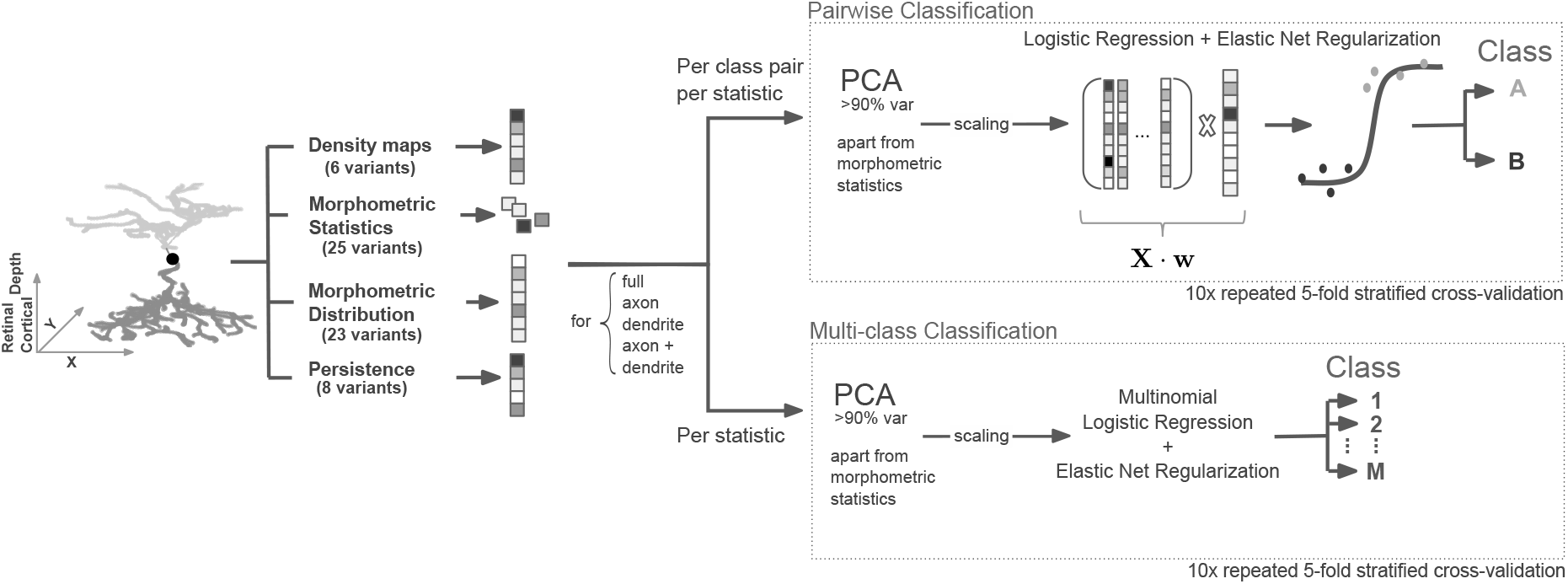
Processing pipeline. Inhibitory interneurons were soma-centered. Retinal bipolar cells were soma-centered in *x* and *y* while *z* = 0 was chosen to correspond to the inner plexiform layer (IPL) onset. The *z* direction of each cell was aligned with cortical/retinal depth, whereas the *x* and *y* direction were left unchanged. Several different feature representations were extracted automatically and used for pairwise and multi-class classifications using logistic regression regularized with elastic net. The performance was assessed using 10 times repeated 5-fold stratified cross-validation.

As an example, Figure 4 shows the performance of one particular feature representation (XZ density map of the full neuron) for all 55 pairs of neural types in the bipolar data set, 21 pairs in the V1 L2/3 data set, 21 pairs in the V1 L4 data set, and 15 pairs in the V1 L5 data set. We used the cross-validated log-loss as the main measure of performance, because it is a proper scoring rule used by logistic regression, it is unaffected by class imbalance and it penalizes confident but wrong decisions. Zero loss means perfect classification, while chance-level performance (for balanced classes) corresponds to the loss of ln(2) ≈ 0.69. For each pair of types, we also computed cross-validated classification accuracy, F1 score, and Matthews correlation coefficient. In our data, the relationships between log-loss and these other performance measures were monotonic and approximately quadratic (Figure S1). As a rule of thumb, a log-loss of 0.2 roughly corresponded to 95% accuracy, a log-loss of 0.4 corresponded to 80% accuracy, and a log-loss of 0.6 corresponded to 65% accuracy.

**Figure 4:**
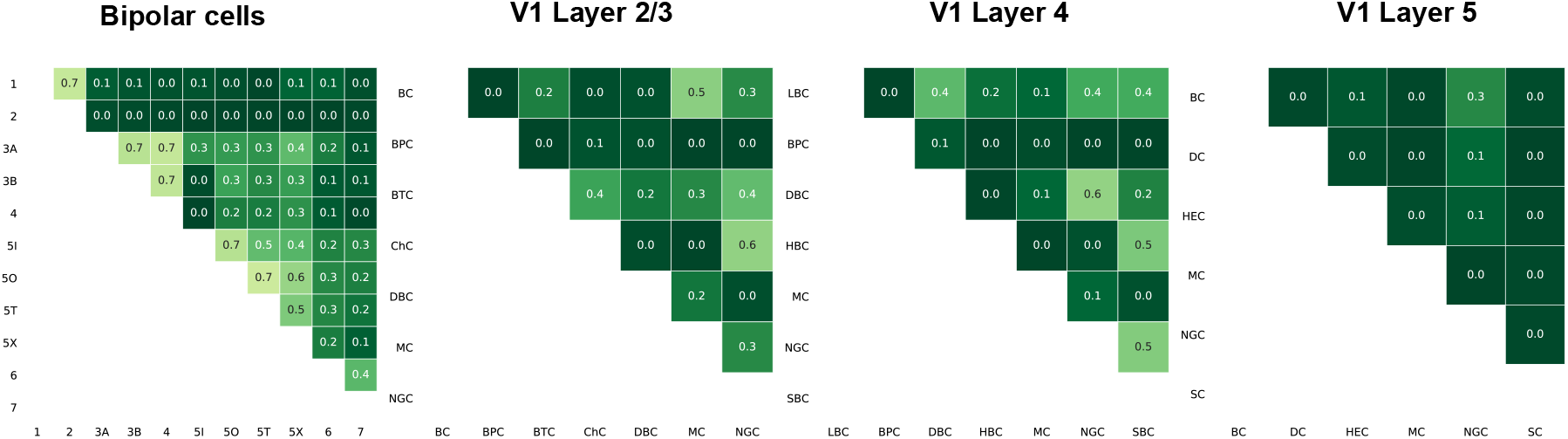
Cross-validated log-loss for each pair of morphological types in each data set using XZ density maps as predictors in logistic regression. Zero log-loss corresponds to perfect prediction, ln(2) *≈* 0.69 corresponds to random guessing. For abbreviations see Figure 1.

The matrix of pairwise classification performances for the bipolar data set can serve as a sanity check that our classification pipeline works as intended: bipolar types with close numbers (e.g. types 1 and 2, or types 3A and 3B) are hard to distinguish (Figure 1), and indeed the log-loss values were generally higher close to the diagonal than far away from it (Figure 4). In fact, to distinguish between bipolar types 1/2, 3A/3B/4, and 5I/5O/5T, the original studies used tiling of the retina and synaptic input patterns in addition to the morphological information (Helmstaedter et al., 2013; Behrens et al., 2016).

For each feature representation, we averaged the log-losses across all pairs within each data set and within each ‘modality’ (full-neuron/axon/dendrite), obtaining 4 × 3 = 12 average log-losses for each of the 62 feature representations. Figure 5 shows a summary for the seven top performing features (see Methods for how they were selected). The performance using the dendritic features was consistently poor for the bipolar cells and the V1 L5 interneurons (close to chance level) and generally much lower than using the axonal features (see also Figure S2). Indeed, for cortical interneurons as well as for retinal bipolar cells, it is the axonal, and not the dendritic, geometry that primarily drives the definition of cell types (Helmstaedter et al., 2013; Sümbül et al., 2014; Ascoli et al., 2008; Markram et al., 2004; DeFelipe et al., 2013), as can be seen in Figure 1. In turn, the performance using the axonal features was practically indistinguishable from the performance of the full-neuron features, consistent with the statistics of our data, where axonal neurites make up about 86% of the total traced neuritic length (4.55 m out of 5.27 m).

**Figure 5:**
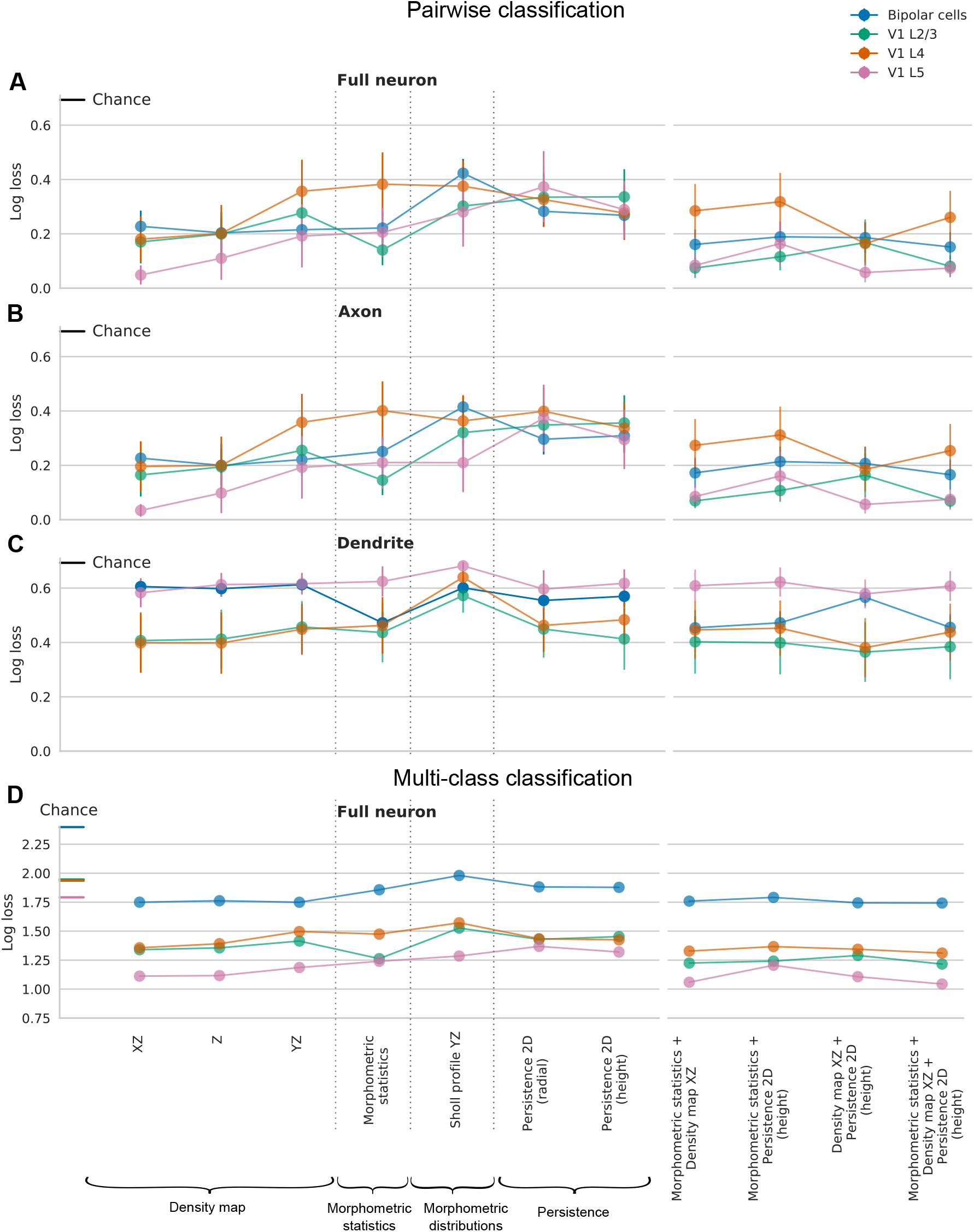
(A–C) Pairwise classification performance of the top performing feature representations based on the full-neuron (A), axonal (B), and dendritic (C) features for each data set. Feature representations are grouped into density maps, morphometric statistics, morphometric distributions, persistence images, and combinations of the top three feature representations. Each shown value is cross-validated log-loss, averaged across all pairs. Error bars correspond to 95% confidence intervals. Chance-level log-loss equals ln(2) *≈* 0.69 and is indicated in each panel. See Figure S2A for the results using combined axonal+dendritic feature representations. **(D)** Cross-validated log-loss of multinomial classification. Chance level for each data set is indicated on the *y*-axis. See Figure S2B for the results using axonal, dendritic, and combined axonal+dendritic feature representations.

Using the performance of the full-neuron features (Figure 5A), we found that the top performing feature representation were XZ density maps, with the mean log-loss of 0.18 ± 0.20 (mean±SD across *n* = 4 datasets), followed by Z density maps (0.19 ± 0.22), morphometric statistics (0.23 ± 0.22), and YZ density maps (0.25 ± 0.23), and then by the 2D persistence images constructed using height (0.29 ± 0.22) and radial distance (0.31 ± 0.22) as filter functions. The best performing morphometric distribution was the YZ Sholl intersection profile (0.37 ± 0.22).

To make a statistical comparison of the performance between two different features *A* and *B*, we computed the mean difference *δ*(*A, B*) in log-loss across all 112 pairs of neural types (pooling pairs across the four data sets). The standard error of *δ* cannot be estimated directly because the pairs are not independent: e.g. the discriminative performances for bipolar types 1 and 2 and for bipolar types 1 and 3A include the same cells from type 1. We used a jackknife procedure *across types* (not across pairs) to estimate the standard error of each reported *δ* (see Methods). We found no evidence of significant difference in performance between XZ density maps and morphometric statistics (*δ* = 0.05 ± 0.05, *z* = 0.92, *p* = 0.36, *z*-test) nor between morphometric statistics and 2D height-based persistence (*δ* = 0.08 ± 0.17, *z* = 0.5, *p* = 0.62). However, the difference between XZ density maps and 2D height-based persistence was statistically significant (*δ* = 0.1 ± 0.04, *z* = 2.7, *p* = 0.007). Among the density maps, the Z density map did not show a significant difference from XZ (*δ* = 0.006 ± 0.02, *z* = 0.37, *p* = 0.71), while YZ was much worse than XZ in the three V1 data sets (*δ* = 0.14±0.05, *z* = 2.84, *p* = 0.005, average across V1 pairs only) but very similar in the bipolar data set (*δ* = 0.01±0.02, *z* = 0.77, *p* = 0.44). Indeed, the *y* direction is mostly meaningless in the V1 data as the slices are flattened during the biocytin staining process (Farhoodi et al., 2019).

For completeness, we also performed all the classifications using axonal and dendritic features pooled together, but the resulting performance was very similar to the performance using axonal (or full-neuron) features alone (Figure S2A).

Next, we asked if combining the feature representations can improve performance. We pooled morphometric statistics and XZ density maps, morphometric statistics and 2D persistence, XZ density maps and 2D persistence, and all three of these feature sets, yielding four additional combined feature sets (Figure 5, right). For some of the data sets (bipolar cells and V1 L2/3) this improved the performance but on average across all data sets we found no increase in performance for feature combinations compared to their constituent feature representations. In some cases performance even decreased, indicating over-fitting due to the small sample sizes in our data (median class size *n* = 16).

### 2.3 Top performing features are consistent across classification schemes

As an alternative to the pairwise classification approach, we also used multi-class classification. We used multinomial logistic regression with exactly the same pipeline of regularization and cross-validation as above. For each of the full-neuron feature representations and each of the data sets, we obtained cross-validated multi-class log-loss (Figure 5D; see Figure S2B for the axonal, dendritic, and combined axonal+dendritic feature representations). Note that for each feature representation and data set, the performance is given by one single estimate, as opposed to the mean over all pairs that we reported above. Therefore only point estimates and no confidence intervals are shown in Figure 5D. Note also that the values of multi-class loss are not directly comparable between data sets, because they are strongly influenced by the number of classes in a data set (*K*). The chance-level performance is given by ln(*K*) and is therefore different for each data set: ln(11) ≈ 2.40 for the bipolar data set, ln(7) ≈ 1.95 for the V1 L2/3 and L4 data sets, and ln(6) ≈ 1.79 for the V1 L5 data set. For this reason, here we are not reporting averages across data sets.

The overall pattern was in good qualitative agreement with that obtained using pairwise classifications (Figure 5D). For three out of four data sets (bipolar, V1 L4 and V1 L5), XZ density maps performed the best (bipolar data set log-loss: 1.75, V1 L4: 1.36, V1 L5: 1.11), followed by morphometric statistics (1.86/1.47/1.24). For the fourth data set (V1 L2/3), morphometric statistics showed the smallest loss (1.26), very closely followed by the density maps (1.34). Combining morphometric statistics with XZ density maps led to a clear improvement in all cortical data sets and was on par with further adding 2D persistence. For the bipolar data, combining features did not improve performance compared to XZ density maps alone. Unlike what we saw above with the pairwise classification, here feature combinations never decreased the performance, possibly due the larger sample sizes of the multi-class classification problems.

To make sure that our conclusions were not dependent on the choice of the classification approach or the performance metric, we repeated the experiments using two other pairwise classifiers: *k*-nearest neighbour (with *k* = 3) and decision trees. In each case we used classification accuracy, F1 score and Matthews correlation coefficient (MCC) as a performance metric (note that log-loss is not available for these classifiers). The selection of top performing feature representations was very consistent, with XZ and Z density maps always ranked the first (Figure 6).

**Figure 6:**
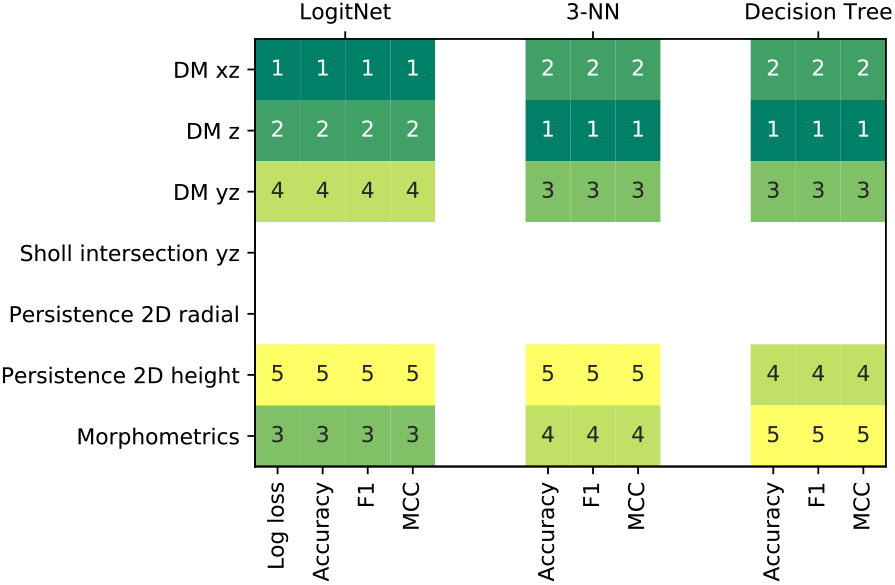
Ranked top five feature representations for each classification scheme using different performance measures on full neuron data. In our case any measure selected the same features within each classification scheme.

### 2.4 The best feature representations are robust against partial tracings

Accurate morphological reconstructions often become more and more difficult to achieve as one goes away from the cell soma, because the neurites become thinner and might have weaker staining which makes them easier to miss. We therefore assessed the robustness of using XZ density maps, morphometric statistics, and 2D persistence as predictors of cell type when neurons are only partially traced.

Partial tracings were simulated by subsequently removing 10 to 90% of the branches (in steps of 10%) of each reconstructed skeleton. On each truncation step, we removed the given fraction of branches with the highest branching order (see Methods). The branching order corresponds to the count of branch points that are passed when tracking the branch back to the soma, so the higher the branching order the more branching has occurred along this branch. This procedure cuts away most of the axonal neurites before reaching the dendrites that typically have branches of lower branch order, and therefore mimics what can happen in actual reconstructions. We used the V1 L2/3 data set for this analysis, performing all pairwise classifications between all pairs of cell types at each truncation step (Figure 7A). In addition, we shuffled the labels of each pairwise comparison to estimate the chance-level distribution of log-losses (Figure 7A, grey shading). Exactly the same cross-validation pipeline was run after shuffling the labels.

**Figure 7:**
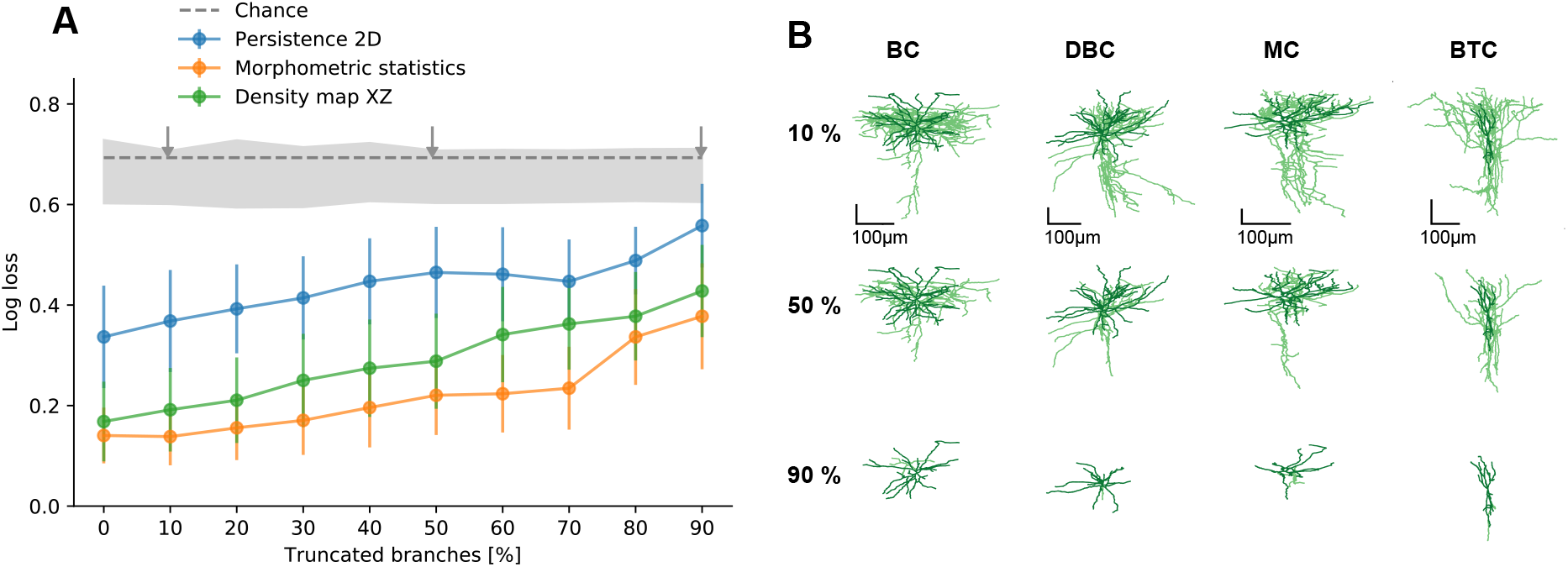
(A) Cross-validated log-loss of XZ density maps, morphometric statistics and 2D persistence (height) as a function of truncation level. Branches were truncated to mimic what happens when neurons are only partially traced. The classification was performed on all pairs of types in V1 L2/3 data set. Dots and error bars show the means and 95% confidence intervals across all 21 pairs. Dashed grey line shows chance level at ln(2) *≈* 0.69. Grey shading shows the chance-level distribution of log-losses obtained by shuffling the labels during the cross-validation (shading intervals go from the minimum to the maximum obtained chance-level values). The arrows mark the levels of truncation shown in panel B. **(B)** XZ projections of four exemplary cells at three levels of truncation: 10%, 50%, and 90%. At 50% truncation the global structure of each cell is still preserved, whereas at 90% only the dendritic structures remain. See Figure 1 for abbreviations.

As expected, performance of each feature representation gradually decreased with increasing level of truncation. The decrease was rather moderate until around 30% truncation level (e.g. it grew from 0.14 ± 0.17 to 0.17 ± 0.19, mean±SD across all 21 pairs, for morphometric statistics, and very similarly for the density map). After that, all representations were noticeably losing in performance.

### 2.5 Using morphological features for unsupervised learning

So far we used supervised learning and assumed cell type labels to be known. A more difficult and arguably more interesting task is to identify morphological cell types using unsupervised clustering (Sümbül et al., 2014; Gouwens et al., 2018). The small sample sizes of our data sets make it very challenging to obtain reliable clustering and to compare the clustering performance of various feature representations. Instead, we directly used the best performing feature representations identified above and performed unsupervised dimensionality reduction using t-distributed stochastic neighbour embedding (t-SNE) (van der Maaten and Hinton, 2008). If the cell types are well-separated in the t-SNE embedding, then it is plausible that a clustering algorithm would identify them as separate types, given a large enough data set.

We first used XZ density maps (reduced to a set of 6–18 PCs capturing 90% of the variance) as an input to t-SNE with perplexity 50 (Figure 8A). The resulting embeddings corresponded well to the pairwise classification performance for XZ density maps that we presented earlier (Figure 4). For example, horizontally elongated cells (HECs) and shrub cells (SCs) in the V1 L5 data set that were both easily distinguishable from other types in the classification task, formed clear clusters, away from other cell types. In contrast, the embedding for basket cells (BCs) and neurogliaform cells (NGCs), the only cell pair with a high log-loss for density maps, showed some overlap. Similarly, retinal bipolar types that were hard to classify, such as types 1 and 2 or types 3A, 3B, and 4, formed joint clusters with a lot of overlap (Figure 8A).

**Figure 8:**
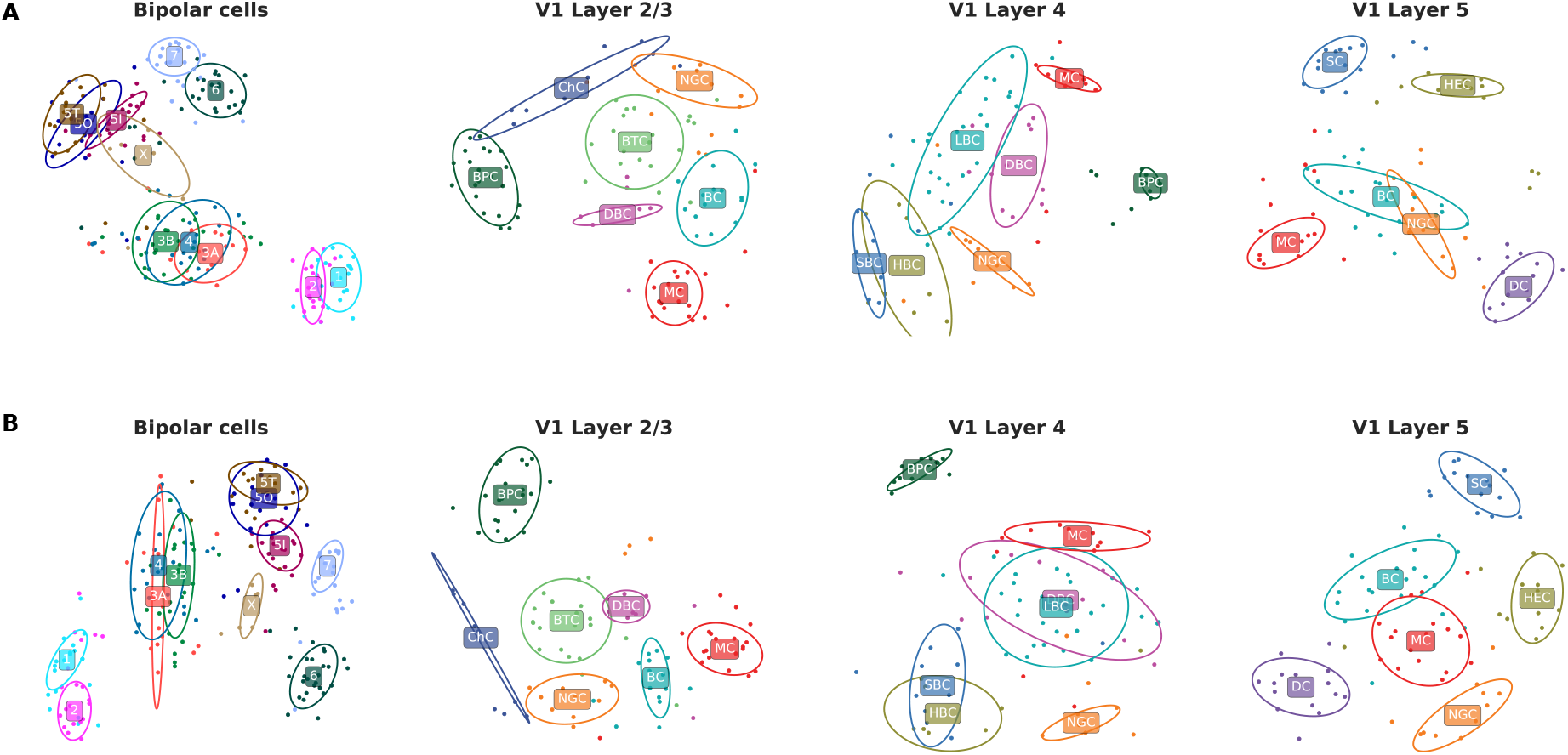
(A) T-SNE embeddings of all four data sets using full-neuron XZ density maps. **(B)** T-SNE embeddings using XZ density maps combined with morphometric statistics. The ellipses are 95% coverage ellipses for each type, assuming Gaussian distribution and using robust estimates of location and covariance. They are not influenced by single outliers. For abbreviations see Figure 1.

We then combined XZ density maps with morphometric statistics. To do so, we reduced each feature representation to a set of PCs capturing 90% of the variance (6–18 PCs) and normalized each set of PCs by the standard deviation of the respective PC1, to put both sets roughly on the same scale. We pooled the scaled PCs together and used this feature representation as input for t-SNE (Figure 8B). For some of the data sets (e.g. V1 L2/3 and V1 L5), the combination of morphometric statistics and XZ density maps yielded an arguably superior t-SNE embeddings with less overlap between types.

The embeddings shown in Figure 8 used full-neuron features that, as we saw above, are dominated by the axonal geometry. Applying the same procedure to the dendritic features yielded embeddings with far worse separation between cell types (Figure S4). Moreover, dendritic features resulted in t-SNE embeddings with far less structure than the full-neuron features, suggesting that there is less of an interesting variability in the dendritic morphologies compared to the axonal ones.

## 3 Discussion

Here we benchmarked existing morphological representations in the context of supervised cell type classification on well-curated data sets encompassing over 500 full reconstructions of interneurons in the mouse visual system. We found that density maps yield the best predictions of cell type labels, followed by morphometric statistics and then 2D persistence images, and showed that they do so even if substantial parts of the traced morphologies are removed. We demonstrated that these predictors work well independent of the used classification scheme or the performance metric suggesting that they are a good starting point for morphological analysis.

### Previous literature

Previous literature has argued that on their own, density maps, morphometric statistics, and persistence work well for cell type identification and classification. Retinal cells, for example, can be successfully discriminated by their stratification depth within the inner plexiform layer (IPL) which can be seen as a *z*-projection of their neurite density (Helmstaedter et al., 2013; Sümbül et al., 2014). Morphometric statistics have been used in a wide variety of studies across species and brain areas (Lu et al., 2013; Polavaram et al., 2014; Lu et al., 2015; Gouwens et al., 2018) and have been shown to perform well in a one-vs-rest classification of cortical neurons (Mihaljević et al., 2015, 2018). Persistence, in turn, has lately been shown to distinguish pyramidal neuron types in juvenile rat somatosensory cortex (Kanari et al., 2019).

These studies, however, did not directly compare different morphology representations, but rather focused on comparing classification schemes or establishing cell type related differences within their chosen morphological representation. Our study fills this gap by applying the same standardized classification procedure to each morphological representation, using well-curated data sets with well-defined cell types. This comparison revealed that the density maps contain enough information to accurately discriminate most inhibitory cell types. This implies that the spatial extent and overall shape of the axonal arbour, as a consequence of a neuron’s connectivity, are more relevant than precise branching characteristics; a finding that has already been proposed for dendrites (Cuntz et al., 2007, 2008, 2010; Cuntz, 2012; van Pelt and van Ooyen, 2013).

Morphometric statistics performed similarly well on some of the data sets but showed a somewhat lower performance than density maps. For example, they failed to distinguish double-bouquet cells and Martinotti cells in layer 4. Various forms of persistence-based measures performed consistently worse than density maps, at least for the interneurons studied here (originally, persistence diagrams were developed for cortical pyramidal neurons (Adams et al., 2017; Kanari et al., 2019; Li et al., 2017)). We did not extensively evaluate combinations of feature representations, as combining even the best representations did not dramatically improve performance, possible as a result of overfitting to a data set of limited size (see Figure 5).

The axonal morphology of neurons in our study contained more information about the cell type than the dendritic morphology, in agreement with the existing literature (Mihaljević et al., 2015; Jiang et al., 2015; Ofer et al., 2018). Our unsupervised analysis also demonstrated far more variability within axonal features compared to the dendritic features (Figure S4), which is in line with classical expert-based cell type naming conventions (Markram et al., 2004; DeFelipe et al., 2013; Helmstaedter et al., 2013). Notwithstanding, dendritic reconstructions are more prevalent in the literature and in the available databases: at the time of writing only 55% (367/667) and 8% (630/8 129) of mouse cortical neurons in the Allen Cell Type atlas (http://celltypes.brain-map.org/data) and the NeuroMorpho library (Ascoli et al., 2007) are flagged as containing complete axonal reconstructions. This is because dendrites are usually thicker and more compact than axons and so are easier to stain and trace. This has obstructed acquisition of complete axonal reconstructions in mammals but might be remedied by recently developed whole brain imaging techniques (Ragan et al., 2012; Yuan et al., 2015; Economo et al., 2016; Gong et al., 2016). At the same time, the robust classification performance of truncated morphologies and the good performance of density maps suggest that full reconstructions might not be necessary for morphological cell type identification.

### Limitations

First, we always used the cell type labels provided in the original publications, treating them as ground truth. For cortical interneurons it has been shown that there is a considerable inter-expert variability between assigned cell type labels (DeFelipe et al., 2013), affecting the outcome of any supervised learning task. Ideally, one would use consensus labels between multiple experts or data modalities for a benchmark evaluation, but such data sets are currently even harder to obtain than the data sets used in this study.

Next, tissue shrinkage and staining method can affect the measured morphology (Farhoodi et al., 2019). In our study all three data sets obtained through biocytin staining (V1 L2/3, V1 L4, V1 L5) showed flattening of the cortical slice (*y*-direction) which made XY density maps perform worse in comparison to other projections. We did not observe this effect for the bipolar cells that have been obtained through EM imaging. Obviously, a feature representation can only be good for classification, if the data contains the relevant information in the first place. Therefore, it is important to be aware of biases or distortions in the experimental protocol before deciding on which feature representation to use.

Further, we believe that a meaningful comparison between different numerical descriptions of morphology is only possible through maintaining strict data consistence and quality criteria. This is why we restricted this work to data from only one species (mouse) of one developmental stage (adult) where morphological cell types are well established and supported by other studies based on electrophysiology (Jiang et al., 2015) or genetics (Shekhar et al., 2016). At the same time we wanted morphologies to be similar enough to pose a challenge and we required complete axonal and dendritic reconstructions. The resulting data set of 505 interneurons is comparable to the sample sizes used in related studies (Mihaljević et al., 2015, 2018). As these interneurons are only locally projecting, our study does not provide guidance as to which features are useful for discriminating neurons based on their long-range projection patterns (Costa et al., 2016; Economo et al., 2018; Gerfen et al., 2018).

As the 505 neurons were split into 31 types, the median sample size per class was only 16 cells. It is difficult to fit machine learning algorithms in the *n* ≪ *p* regime where the number of dimensions *p* highly exceeds the number of samples *n* (Friedman et al., 2001). We used a simple linear model strongly regularized by PCA preprocessing and an elastic net penalty as well as nonlinear non parametric models since fitting more complicated models be challenging with these low sample sizes. This approach performs well when the leading principal components of the data have good discriminative power, but can also perform at chance level if the difference between types is restricted to low-variance directions. Thus, low classification performance for a given cell type pair does not necessarily imply that they could not be reliably separated with more available data.

Finally, we restricted our benchmarking effort to the most prominent and established morphological representations that have been independently employed by more than one research group. In particular, this excluded some methods based on graph theory (Heumann and Wittum, 2009; Gillette and Grefenstette, 2009) and sequence alignment (Gillette and Ascoli, 2015; Gillette et al., 2015; Costa et al., 2016) which can be promising candidates for further studies. The morphometric statistics that we used did not include everything that has been suggested in the literature either. For example, we did not use morphometric statistics such as fractal dimension because of their disputed relevance for our data (Panico and Sterling, 1995) and did not explicitly quantify the amount of layer-specific arborization (DeFelipe et al., 2013; Gouwens et al., 2018) because this concept only applies to cortical neurons and layer boundaries were not available for our data. However, given the superior performance of density maps, it is possible that including layer-specific information could improve the performance of morphometric statistics.

### Outlook

Our study serves to provide a starting point for future work on algorithmic cell type discrimination based on anatomical data, for example in the context of large-scale efforts to map every cell type in the brain as pursued by the NIH BRAIN initiative. It allows experimenters to make an informed choice which cell type representations are useful to automatically distinguish interneurons based on their morphology. Of course, how far our results generalize to other species remains to be seen. The resulting representations also make it possible to relate anatomical descriptions of neurons to data from other modalities such as e.g. gene expression patterns (Cadwell et al., 2017).

The representations investigated here are purely descriptive and do not provide deeper mechanistic insight, compared e.g. to generative models of the growth process of neurons during development (van Pelt and Schierwagen, 2004; Cuntz et al., 2010; Memelli et al., 2013; Wolf et al., 2013; Fard et al., 2018; Farhoodi et al., 2019). Ideally, a mechanistically grounded feature representation would perform at least on par with density maps for cell type discrimination while yielding parameters that are more easily interpretable. Potential starting points for such a representations are growth models proposed by van Pelt and Schierwagen (2004) and Cuntz et al. (2010), which have a manageable amount of parameters and show systematic parameter differences for dendrites of different cell types. This make them promising candidates for further research.

## 4 Methods

### 4.1 Data

We used data from Helmstaedter et al. (2013), Scala et al. (2019), and Jiang et al. (2015), splitting the latter data set into two parts by cortical layer. All neurons were labelled by human experts in the original studies. We confirmed the quality of all reconstructions through inspection. Our study investigated a total of 5.27 meters of traced neurites from *n* = 505 neurons.

1. **Bipolar cells.** This data set comprised *n* = 221 tracings of retinal bipolar cells in one mouse (p30) from electron-microscopy data (Helmstaedter et al., 2013). To allow for at least 5-fold cross-validation, we did not analyze cell types which had counts of 5 cells or fewer. This criterion excluded types 8 and 9 and resulted in *n* = 212 remaining morphologies in 11 types. The reconstructions (as .SWC files) as well as their cell type labels were obtained from the authors of Behrens et al. (2016) which explains the additional cell types 5O, 5I and 5T as compared to the original work.
2. **V1 Layer 2/3.** Manually traced biocytin stainings of *n* = 108 inhibitory interneurons of 7 types in layer 2/3 (L2/3) of adult mouse primary visual cortex (Jiang et al., 2015). We obtained the reconstructions (as .ASC file) and their cell type labels from the authors.
3. **V1 Layer 4.** Manually traced biocytin stainings of *n* = 92 inhibitory interneurons of 7 types in layer 4 (L4) of adult mouse primary visual cortex (Scala et al., 2019). We obtained the reconstructions (as .ASC file) and their cell type labels from the authors.
4. **V1 Layer 5.** Manually traced biocytin stainings of *n* = 94 inhibitory interneurons of 6 types in layer 5 (L5) of adult mouse primary visual cortex (Jiang et al., 2015). One deep-projecting cell lacked an axon so it was excluded from further analysis resulting in *n* = 93 remaining morphologies. We obtained the reconstructions (as .ASC file) and their cell type labels from the authors.

For data availability see Information Sharing Statement.

### 4.2 Preprocessing and nomenclature

Reconstructed morphologies were converted into SWC format using `NLMorphologyConverter` 0.9.0 (http://neuronland.org) where needed and further analysed in Python. The SWC format represents a morphology with a list of *nodes* (points) with each node described by its *id*, *3D position*, *radius*, *type* (1: soma, 2: axon, 3: dendrite), and *parent id*. Each node connects to its parent node with a straight line that we will call “sub-segment”. Several nodes can connect to the same parent note; in this case this parent node is called a “branch point”. A neurite path from one branch point to the next is called a “segment”.

The bipolar cells were missing explicit type labels for the soma, we therefore set every node of radius larger than 1 micron to be somatic. We generally allowed for only one somatic node so that within one reconstruction all somatic nodes were grouped and replaced by one node with position and radius being the mean across all original soma nodes. Especially in the initial branch segments it can occur that node type labels (1: soma, 2: axon, 3: dendrite) are not consistent between consecutive nodes. Node type labels within one branch we hence assigned according to the majority vote over all sub-segment types within this branch.

All cortical interneurons were soma-centered and their *z* coordinate (height) was oriented along the cortical depth. We did not account for the shrinkage of slice thickness (depth) that happens during the staining, however, we ameliorated reconstruction jumps in *y* direction using a 3rd order Savitzky-Golay filter with window size of 21 microns (as implemented in `scipy.signal`). For this we re-sampled all neurons to have equidistant points of 1 micron along all neurites. Bipolar cells were soma-centered in their *x* and *y* coordinates and IPL-centered in the *z* direction corresponding to the retinal depth (i.e. *z* = 0 corresponded to the outer border of the inner plexiform layer, IPL). Smoothing and re-sampling was not done for this data set.

### 4.3 Feature representations

We calculated 62 different feature representations for each cell. These representations can be grouped into four categories: *density maps*, *morphometric statistics*, *morphometric distributions*, and *persistence*. All feature representations were separately computed for axons, for dendrites, and for the whole neuron (without distinguishing axons from dendrites) yielding 62 · 3 = 186 representations per neuron (see Figure 2).

### 4.3.1 Density maps

We sampled equidistant points with 25 nm spacing along each neurite of the traced skeletons and normalized the resulting point clouds within each data set for each modality to lie between 0 and 1 (for the range values used for this normalisation see the linked Github repository). For 2D density maps, the normalized point cloud was projected onto the *xy*, *xz*, and *yz* planes and binned into 100 × 100 bins spanning [−0.1, 1.1]. For 1D density maps the normalized point cloud was projected onto the *x*, *y*, and *z* axes and binned into 100 bins spanning [−0.1, 1.1]. We smoothed the resulting histograms by convolving them with a 11 × 11 (for 2D) or 11-bin Gaussian kernel with standard deviation *σ* = 2 bins. For the purposes of downstream analysis, we treated the density maps as vectors of 10 000 (for 2D) or 100 (for 1D) features. Overall we used 6 versions of density map representations.

#### 4.3.2 Morphometric statistics

For each cell we computed a set of 24 single-valued summary statistics:

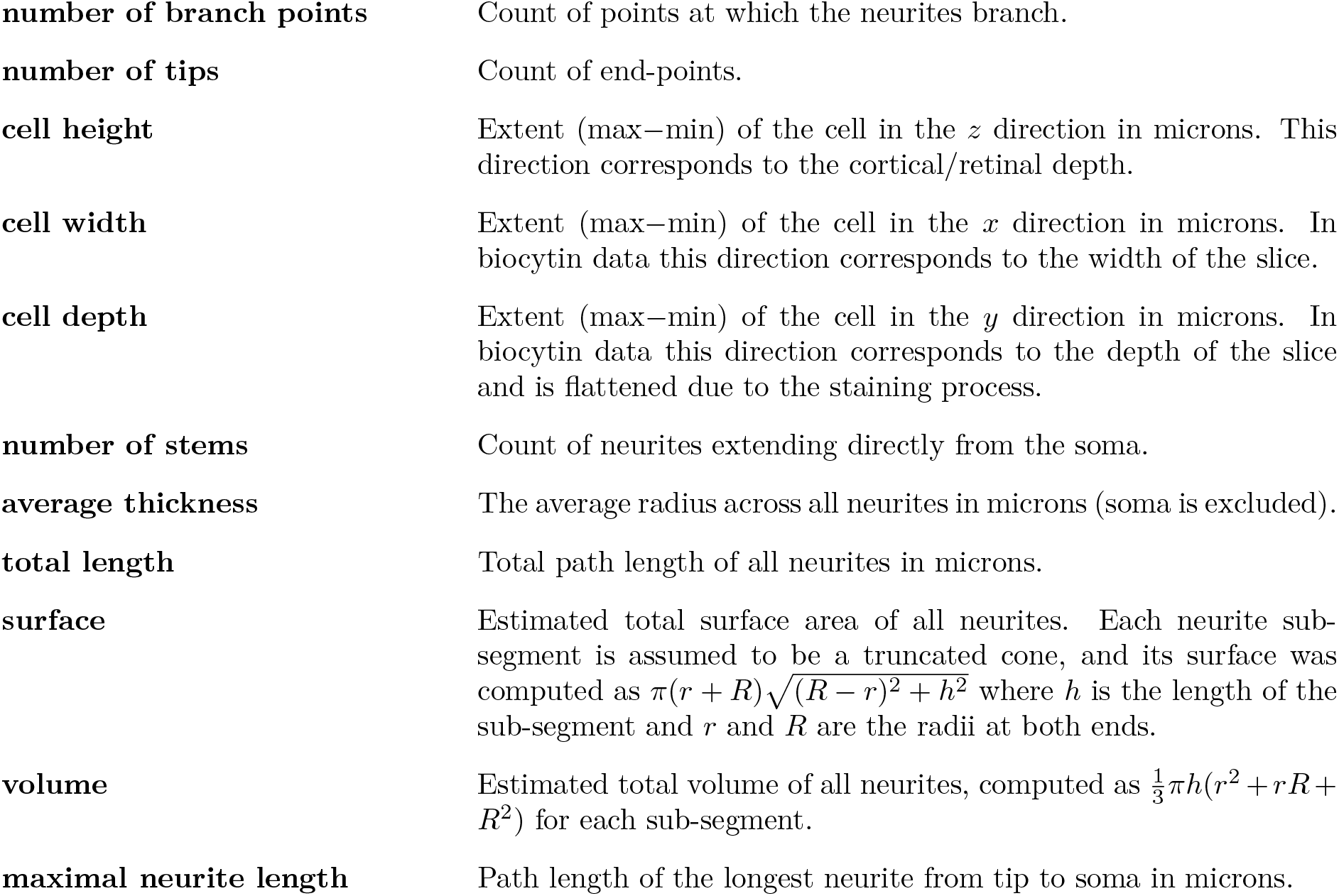
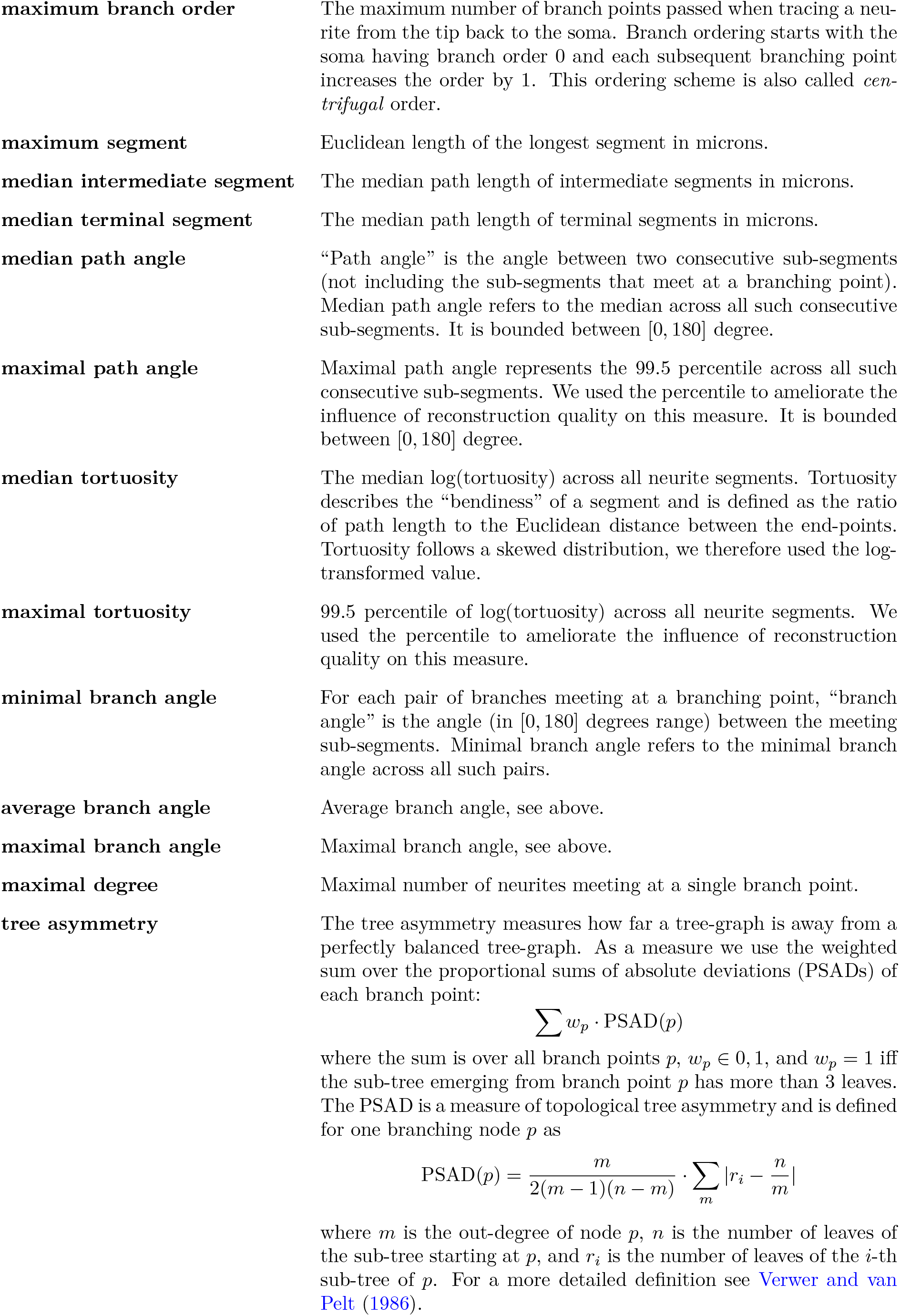

In addition to the 24 features listed above, we grouped all of them together into one vector which we named *morphometric statistics*.

#### 4.3.3 Morphometric distributions

For each cell we computed the following 17 one-dimensional morphometric distributions:

|  |  |
| --- | --- |
| **branch angles** | Histogram of bifurcation angles between neurites (20 bins from 0 to 180). |
| **branch orders** | A vector of length $K$ where $K$ is the maximal branch order within each data set, with element $i$ equal to the number of branch-points of order $i$. |
| **path angles** | Histogram of the path angles (20 bins from 0 to 180). |
| **root angles** | Histogram of the root angles of each segment. The root angle denotes the angle between the straight line connecting the segment's start and end nodes and the straight line that connects the segment's end point with the soma (20 bins from 0 to 180). It is indicative of the preferred growing direction of the neural arbor. For more details see Bird and Cuntz (2019). |
| **Euler root angles, $\alpha$** | Histogram of the Euler root angles ($\alpha$). Here, the root angles are expressed as Euler angles around the $x$, $y$ and $z$ axis (20 bins from 0 to 180). |
| **Euler root angles, $\beta$** | See above. |
| **Euler root angles, $\gamma$** | See above. |
| **segment lengths** | Histogram of Euclidean segment lengths (20 bins from 0 to the maximal segment length within the data set). |
| **thickness** | Histogram of nodes' radii, the soma is excluded (30 bins from 0 to the maximal radius within the data set). |
| **path distance to soma** | Histogram of the path length of each branch point and tip to the soma (20 bins from 0 to maximal neurite length within data set) |
| **Euclidean distances to soma** | Histogram of the Euclidean distance of each branch point and tip to the soma. |
| **Sholl intersection $xy$** | Sholl intersection profile in the $xy$-plane. The Sholl intersection profile describes the number of intersection of a 2D projection with concentric circles of different radius around the soma (Sholl, 1953). We used 36 steps from soma to maximal radial distance from soma. |
| **Sholl intersection $xz$** | Sholl intersection profile in the $xz$-plane. |
| **Sholl intersection $yz$** | Sholl intersection profile in the $yz$-plane. |
| **3-star motif** | 10-dimensional vector with element $i$ being the number of 3-star motifs in the sub-graph containing $i \cdot 10\%$ of all nodes closest to the soma. A 3-star motif represents a branch point where one branch forks into two branches. For each additional branch the 3-star motif is counted again (e.g. a neurite forking into three sub-branches is counted as having two 3-star motifs). |
| **average 3-star motif** | The 3-star motif vector is calculated 100 times using different random nodes instead of the soma as the "center" of concentric spheres. The resulting vectors are then averaged. |
| **average maximal distance** | Using the same sub-graphs, we compute the maximal Euclidean distance in each sub-graph (10-dimensional vector). This procedure is repeated 100 times with random starting nodes and the resulting vectors are then averaged. |

In addition, we used six two-dimensional morphometric distributions. The binning and normalization were the same as for the respective 1D distributions

|  |  |
| --- | --- |
| **branch angles × branch orders** | 2D histogram of branch angles as a function of branch orders (across all branch points). |
| **branch angles × path distances** | 2D histogram of branch angles as a function of path distances to the soma in microns (across all branch points). |
| **path angle × branch order** | 2D histogram of path angles as a function of branch orders (across all nodes, not only branch points). |
| **path angle × path distance** | 2D histogram of path angles as a function of path distances to the soma in microns (across all nodes). |
| **thickness × branch order** | 2D histogram of neurite radii as a function of branch orders (across all nodes). |
| **thickness × path distance** | 2D histogram of neurite radii as a function of path distances to the soma in microns (across all nodes). |

#### 4.3.4 Persistence diagrams

Persistence diagrams originate from the field of algebraic topology but recently have been proposed as a representation for neural morphologies (Kanari et al., 2018). We briefly outline their underlying algorithm here. Starting from each tip, one records the “birth time” of each branch as the distance of the tip from the soma. Hereby, the distance is measured according to some filter function *f*. While moving away from the tips towards the soma, at each branch-point the “younger” branch, i.e. the one with a smaller birth time, is “killed” and its “death time” is recorded as the distance of the branch point from the soma. This results in a 2D point cloud of (*birth time*, *death time*) for each branch, the so called *persistence diagram*, in which only the longest branch survived until the soma and has a death time of 0. Depending on the filter function *f*, different aspects of the neuron’s topology can be captured. Here we employed four different filter functions, following Kanari et al. (2018):

**radial distance** returns the Euclidean distance of node *x* to the soma *s*;

**path length** returns the sum over the lengths of all segments along the path connecting node *x* with the soma *s*;

**branch order** returns the branch order of node *x*;

**height** returns the difference of *z*-coordinates between node *x* and the soma *s* projected onto the *z*-axis.

Euclidean distance is undefined for persistence diagrams themselves. To circumvent this problem we converted each persistence diagram into a 1D or 2D image and used Euclidean distance on the results as this procedure has been shown to work well in the neural domain (Kanari et al., 2019; Li et al., 2017). To obtain a 2D Gaussian persistence image we performed kernel density estimation of the point cloud using a 2D Gaussian kernel (`gaussian_kde` from the `scipy.stats` package with default settings). We evaluated the density estimate on a 100 × 100 equidistant grid spanning a [0, max_birth_] × [0, max_death_] rectangle. Here max_birth_ and max_death_ refer to the maxima across all cells within each data set (for actual values see the linked Github repository). For the purposes of downstream analysis, we treated this as a set of 10, 000 features.

The 1D Gaussian persistence vector we obtained in a similar way. Namely, we performed a one-dimensional Gaussian kernel density estimation of the neurites’ “living time” (birth−death) and sampled the resulting estimate at 100 equidistant points spanning [0, max_birth_].

### 4.4 Classification

Each feature representation was used as a predictor for pairwise and multinomial classification. Except for morphometric statistics, we reduced all representations using principal component analysis (PCA) and kept as many features as needed to capture at least 90% of the variance on the training set (for cross-validation, PCA was computed on each outer-loop training set separately, and the same transformation was applied to the corresponding outer-loop test set). We divided all PCA components by the standard deviation of the respective first principle component to put all features roughly on the same scale and to allow for combination of features. Morphometrics were z-scored.

For binary classification, we used logistic regression with an elastic net regularization. It minimizes the following loss function:

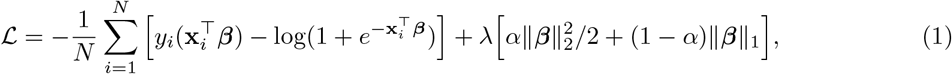

where **x**_*i*_ are predictors, *y_i_* a binary response (that can be 0 or 1), and there are *N* training samples. Regularization parameter *α* was fixed to 0.5, which is giving equal weights to the lasso and ridge penalties. We used nested cross-validation to choose the optimal value of the regularization parameter *λ* and to obtain an unbiased estimate of the performance. The inner loop was performed using the `civisanalytics Python` wrapper around the `glmnet` library (Friedman et al., 2010) that does *K*-fold cross-validation internally (default: 3-fold). We kept the default setting which uses the maximal value of *λ* with cross-validated loss within one standard error of the lowest loss (`lambda_best`) to make the test-set predictions. We explicitly made the `civisanalytics` Python wrapper use the loss (and not accuracy) for *λ* selection:

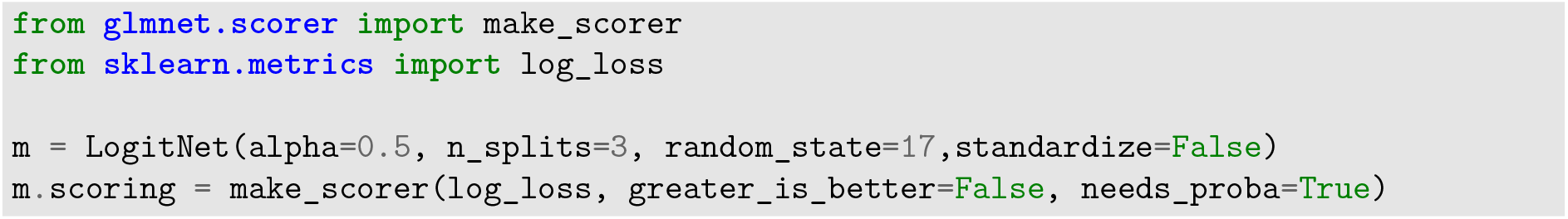

The outer loop was 10 times repeated stratified 5-fold cross-validation, as implemented in `scikit-learn` by

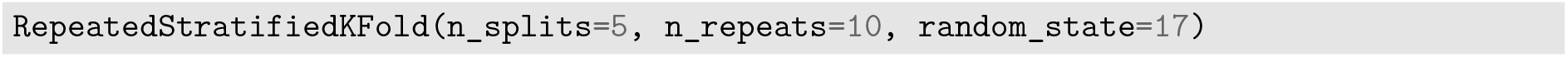

Model performance was assessed via mean test-set log-loss and test-set accuracy. For comparison between different classification schemes we also computed the mean test-set macro F1 scores, which is the unweighted mean of the F1 scores for each class, and the mean test-set Matthews correlation coefficients (Matthews, 1975).

For multi-class classification we used multinomial logistic regression with an elastic net regularization.

The parameters and the cross-validation procedure were the same as above.

The processing pipeline including preprocessing, feature extraction and classification was automated using DataJoint (Yatsenko et al., 2015).

#### 4.4.1 Selection of top performing features

We identified the top five performing feature representations for each “modality” (full-neuron, axon, dendrite, as well as axon + dendrite) based on their mean binary classification performance across data sets and identified their superset (six features). We also included the best performing morphometric distribution to have at least one feature representation of each category investigated. This lead to the seven features shown in Figure 5.

#### 4.4.2 Statistical analysis of differences

We estimated the mean difference between two feature representations *A* and *B* as

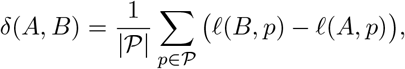

where 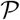 denotes the set of pairs of types across the four data sets, 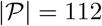 is their total number, and *l*(*X, p*) is cross-validated log loss of feature *X* for pair *p*. To estimate the standard error of *δ*(*A, B*) we use the jackknife procedure (Efron and Hastie, 2016). We repeat the procedure leaving out one type *τ* entirely, so that all pairs including that type are left out:

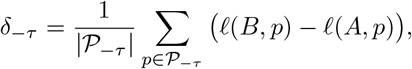

where *P*_−*τ*_ is the set of pairs without type *τ*. This yields *n* = 31 estimates of *δ*_−*τ*_, with the jackknife estimate of the standard error given by

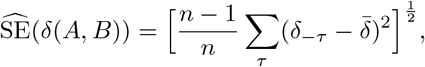

where 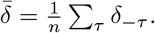. All reported *p*-values were obtained with a *z*-test using

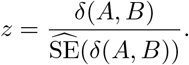

#### 4.4.3 Other classification schemes

For comparison with different classification schemes we used the 3-nearest neighbour classifier and the decision tree classifier of the Python `scikit-learn` implementation:

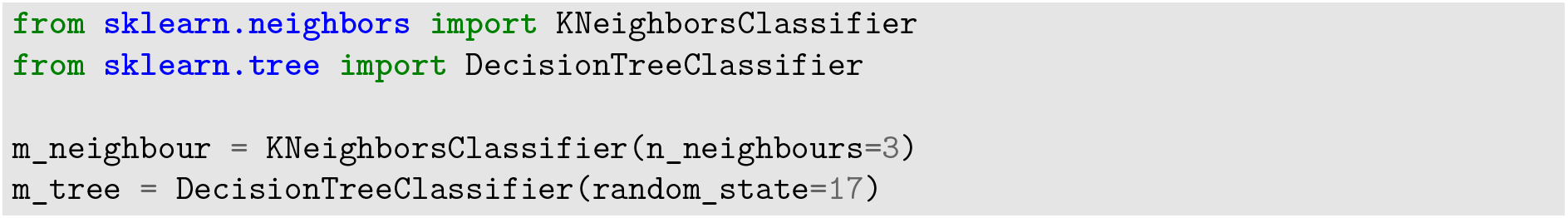

### 4.5 t-SNE visualization

For the t-SNE visualization (van der Maaten and Hinton, 2008) of each data set, we reduced density maps and morphometric statistics to as many principal components needed to keep 90% of the variance each. As done during classification, we scaled each set of PCs by the standard deviation of the respective PC1, to put both sets roughly on the same scale. Then we stacked them together to obtain a combined representation of each cell. Exact (non-approximate) t-SNE was run with perplexity 50 and random initialization using the `scikit-learn` implementation:

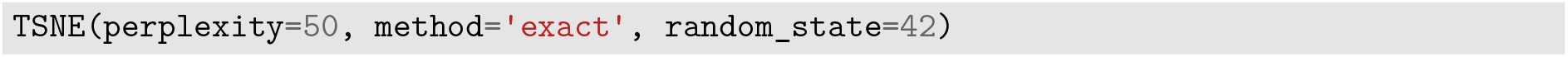

To plot the coverage ellipses for each cell type (Figure 8), we used robust estimates of location and co-variance, so that the ellipses are not influenced by outliers. We used the minimum covariance determinant estimator (Rousseeuw and Driessen, 1999) as implemented in `MinCovDet()` in `scikit-learn`.

### 4.6 Robustness analysis

Morphological tracings are done manually and reconstruction quality can vary between protocols and experts. We assessed the robustness of the top performing feature representations by repeating the classification procedure on only partially traced neurons. We simulated incomplete tracings using the full reconstructions of the V1 L2/3 data set and assessed the performance of all pairwise classifications (Fig. 7). Incomplete tracings were simulated by successively removing 10–90% of all branches starting with the branches of the highest branch order. We then used the XZ density map, the morphometric statistics and the 2D persistence image for each degree of truncation as predictors.

To estimate the chance-level performance we shuffled the class labels for each pairwise classification at each truncation grade and repeated our classification pipeline with shuffled labels. The resulting distribution of chance-level test-set log loss values was in agreement with the theoretical value of ln(2).

### Author contributions

PB and SL designed the study. SL performed the analysis with supervision from DK and PB. All authors wrote the manuscript.

### Information Sharing Statement

Our analysis code is available at: http://github.com/berenslab/morphology-benchmark. The bipolar cell reconstructions will be made available upon publication of the manuscript. The original reconstructions of the V1 interneurons of layer 2/3 and layer 5 are available at http://neuromorpho.org (archive Tolias). Check the “Include Auxiliary files” box to get the originally uploaded files with cortical depth alignment. The morphological reconstructions of V1 layer 4 interneurons(in .asc and .swc formats) are deposited to Zenodo at https://doi.org/10.5281/zenodo.3336165.

## Acknowledgements

We thank Ulrike von Luxburg, Debarghya Ghoshdastidar, Xialong Jiang, Federico Scala and Andreas Tolias for discussions. This work was funded by the German Ministry of Education and Research (FKZ 01GQ1601) and the German Research Foundation (DFG) under Germany’s Excellence Strategy (EXC 2064/1 – 390727645) and grants to PB (BE5601/4-1, SFB 1233 “Robust Vision” - Project number 276693517) and the National Institutes of Health BRAIN Initiative (1U19MH114830-01).

---

**Figure S1:**
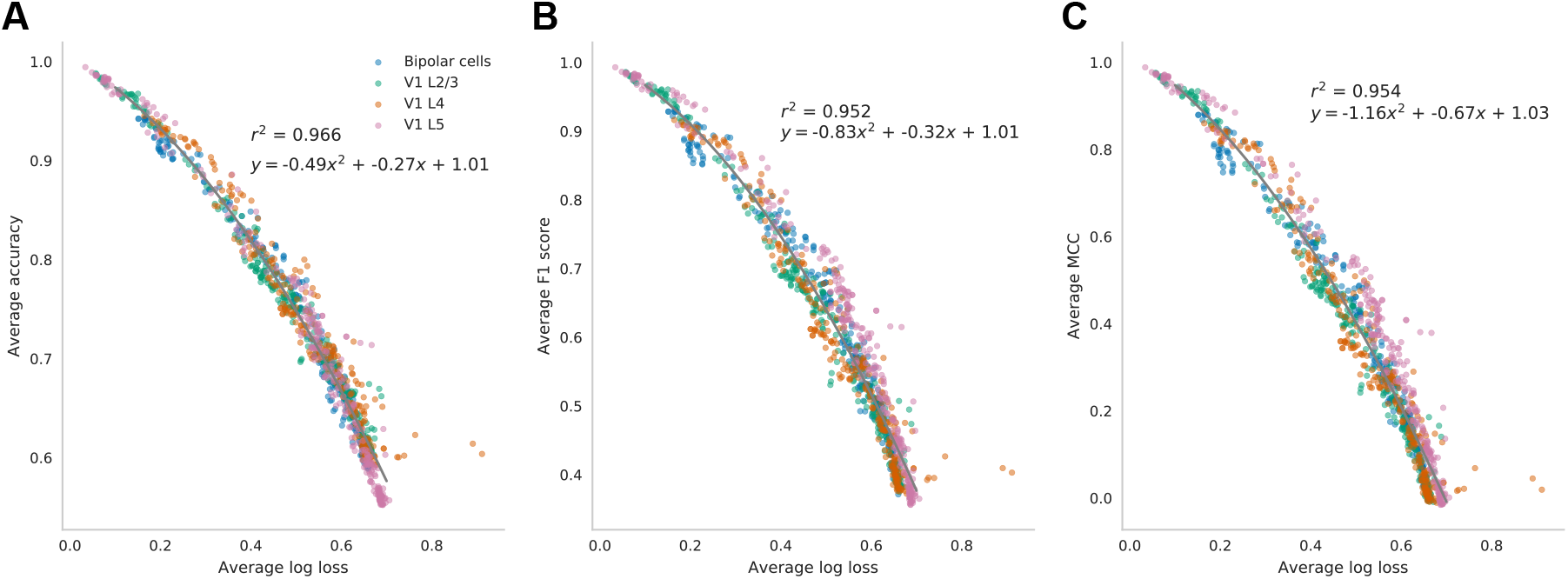
(A–C)Conversion of pairwise classification performance between different average performance metrics. Each dot represents the average across all pairs within one data set for one modality and one selected feature representation. The conversion is shown between average log-loss and average accuracy (**A**), average log-loss and average F1 score (**B**), and average log-loss and average Matthews correlation coefficient (**C**). The grey line denotes a quadratic regression fit.

**Figure S2:**
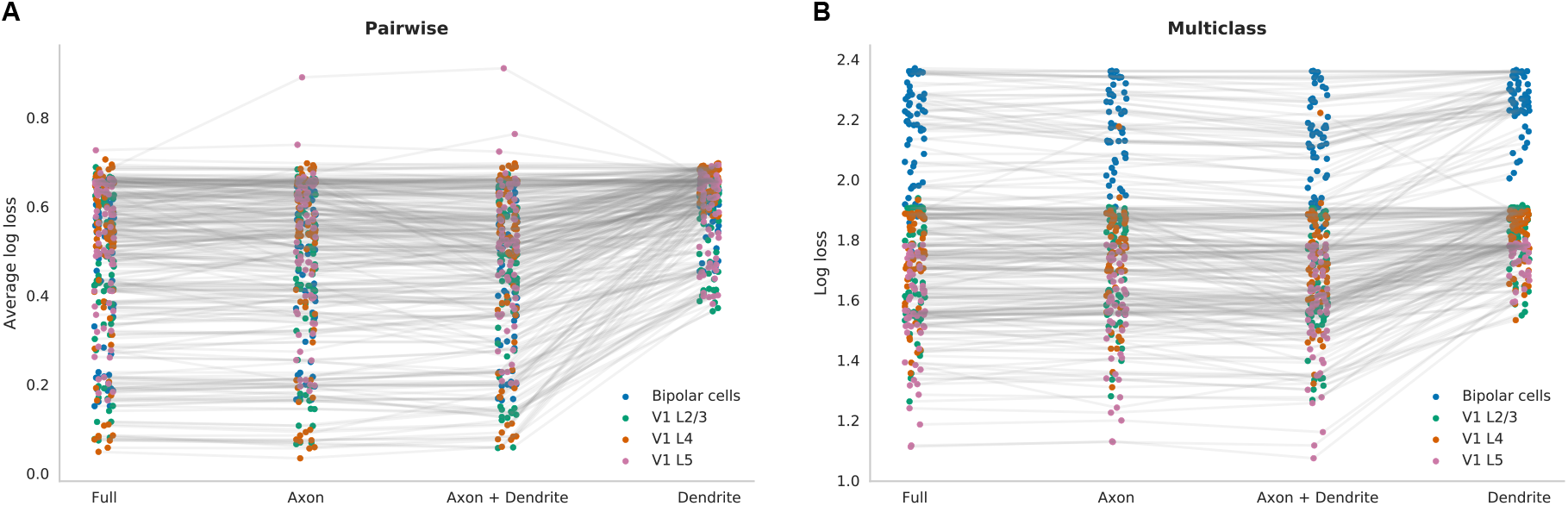
Comparison of average log-losses for each statistic in each modality and data set for binomial (**A**) and for multinomial classification (**B**). In both cases, dendrites perform considerably worse than any of the other modalities.

**Figure S3:**
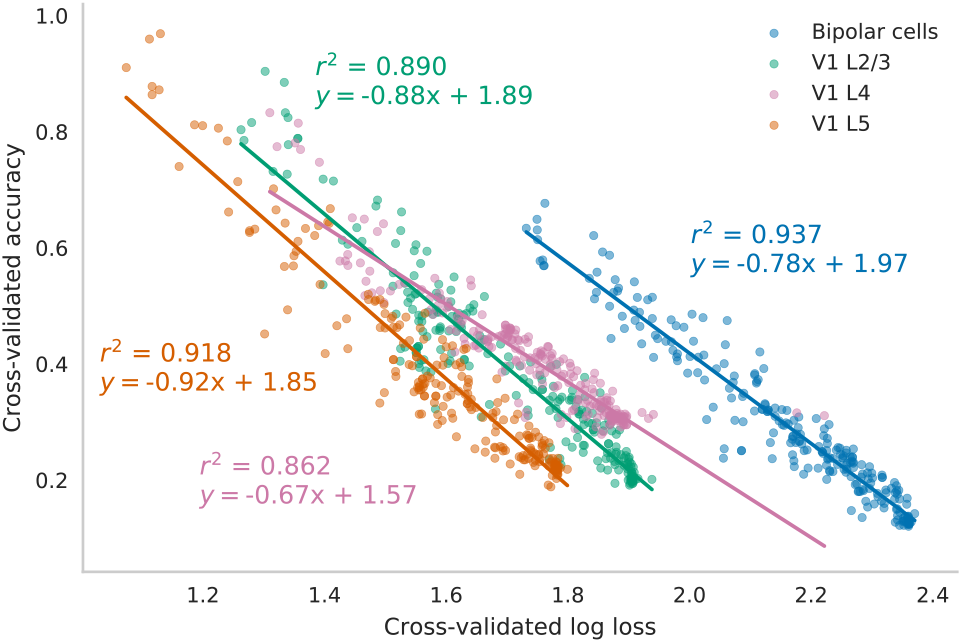
Conversion from multi-class loss to accuracy for each data set. The colored lines denote a linear regression fit to the respective data set. V1 layer 5 (orange) is almost perfectly separable with accuracies close to 1 for some feature representations. This is also indicated by the emerging clusters in Figure 8.

**Figure S4:**
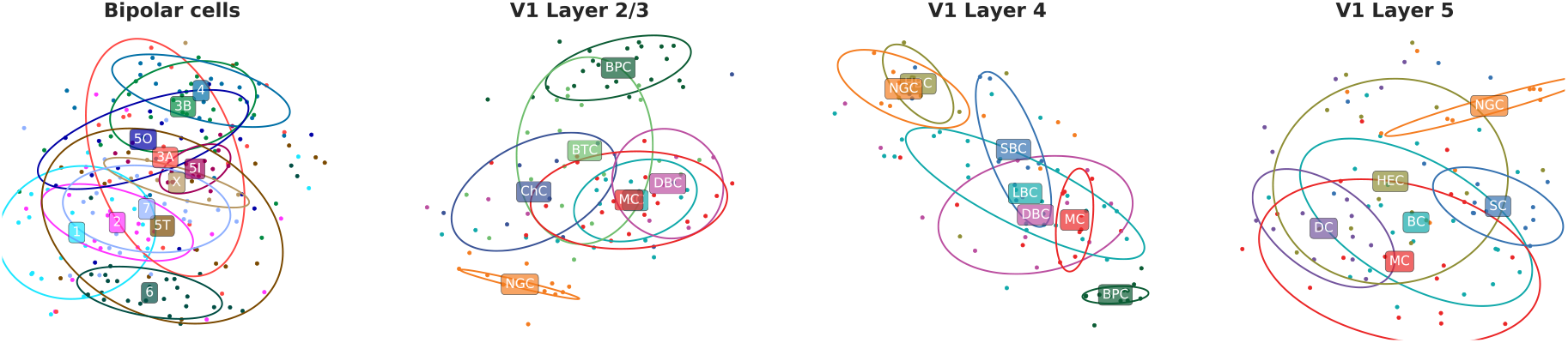
T-SNE embeddings of all four data sets (perplexity 50) for dendrites. We used a combined feature representation consisting of XZ density map and morphometric statistics. Both feature representations were reduced to as many PCs needed to keep 90% of the variance and combined into one feature set prior to t-SNE.

**Figure S5:**
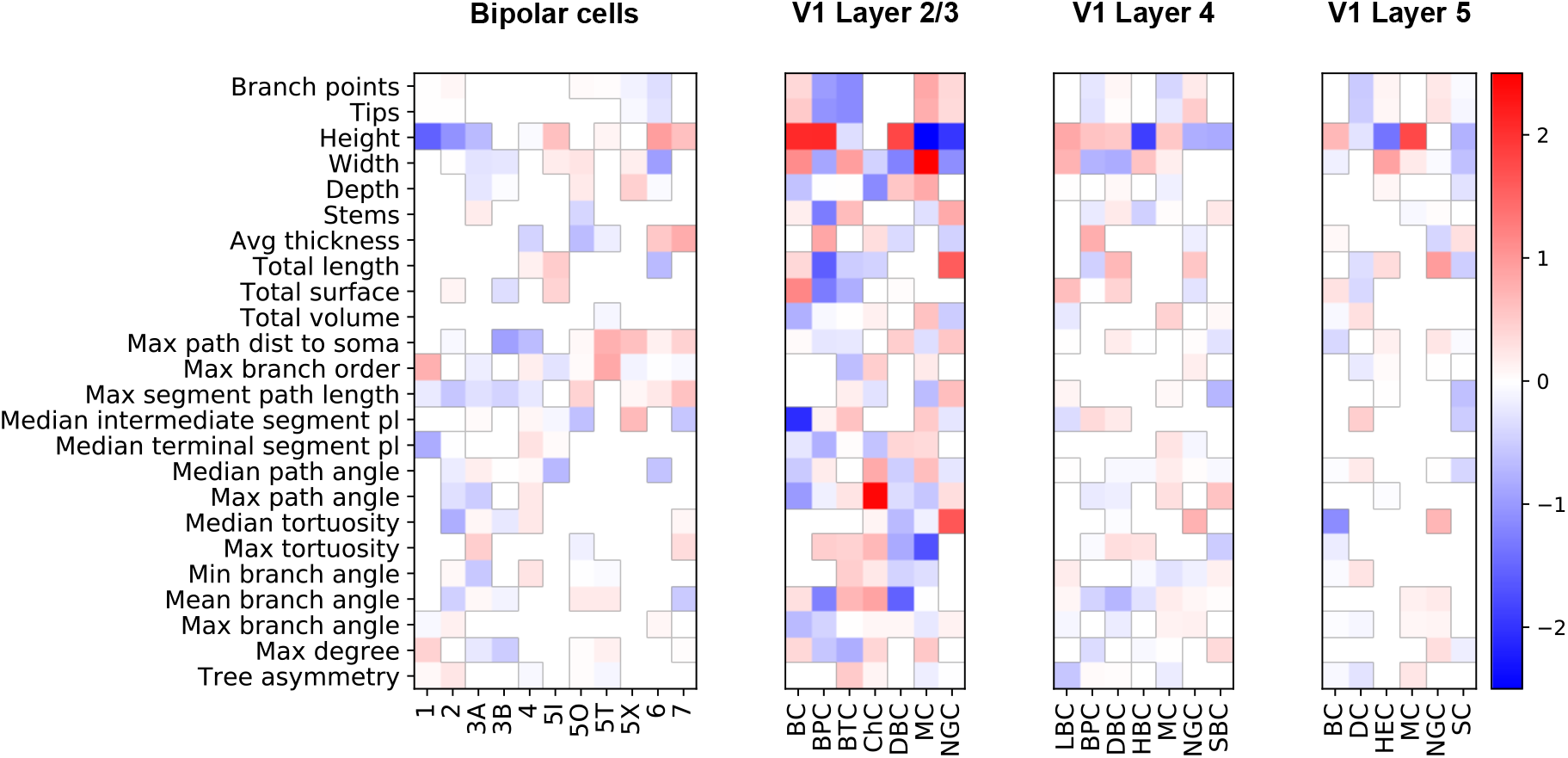
Visualization of the coefficient vectors that are fitted during the multinomial Logistic regression on the morphometric statistics for each data set. The data shown here is based on the full neural reconstructions. White squares indicate a coefficient of zero and have been masked out to improve visibility.

